# Scaffold and structural diversity of the secondary metabolite space of medicinal fungi

**DOI:** 10.1101/2022.09.25.509364

**Authors:** R.P. Vivek-Ananth, Ajaya Kumar Sahoo, Shanmuga Priya Baskaran, Areejit Samal

## Abstract

Medicinal fungi including mushrooms have well documented therapeutic uses. The MeFSAT database provides a curated library of more than 1800 secondary metabolites produced by medicinal fungi for potential use in high throughput screening (HTS) studies. In this study, we perform a cheminformatics based investigation of the scaffold and structural diversity of the secondary metabolite space of medicinal fungi, and moreover, perform a detailed comparison with approved drugs, other natural product libraries and semi-synthetic libraries. We find that the secondary metabolite space of MeFSAT has similar or higher scaffold diversity in comparison to other natural product libraries analysed here. Notably, 94% of the scaffolds in the secondary metabolite space of MeFSAT are not present in the approved drugs. Further, we find that the secondary metabolites of medicinal fungi, on the one hand are structurally far from the approved drugs, while on the other hand are close in terms of molecular properties to approved drugs. Lastly, chemical space visualization using dimensionality reduction methods showed that the secondary metabolite space has minimal overlap with the approved drug space. In a nutshell, our results underscore that the secondary metabolite space of medicinal fungi is a valuable resource for identifying potential lead molecules for natural product based drug discovery.

## Introduction

Natural products, semi-synthetic and synthetic libraries of different sources are being leveraged in high throughput screening (HTS) to identify new anti-virals, anti-bacterials and anti-cancer agents.^1,2^ Moreover, there is an increased focus toward natural product libraries for identification of new chemical entities with immunomodulatory, anti-aging and cognitive enhancement properties to prevent diseases and promote holistic well-being.^3–5^ The selection of appropriate chemical libraries with high diversity for HTS is a critical step in the drug discovery pipeline. Notably, chemical libraries with high structural diversity have a higher hit identification rate in HTS than similarly sized libraries with low structural diversity.^6,7^ Therefore, it is imperative to assess the diversity encoded by the chemical libraries, especially the natural product libraries which are a promising source of diverse chemical scaffolds.

Natural products from plants, fungi, bacteria and marine organisms are rich source of biologically relevant and therapeutic chemical entities.^8^ Several databases of natural products of plants and microbial origin have been developed to facilitate the ongoing efforts in natural product based drug discovery.^9–12^ In spite of the efforts by the scientific community to map the complete natural product space, only a small fraction of the space has been mapped so far. The natural product space of medicinal plants and fungi represents a chemical space wherein the small molecule entities have a higher likelihood of being therapeutic.^9,10,12–14^. Thus, there has been several efforts to develop and analyse phytochemical libraries of medicinal plants used in traditional medicine such as TCM-Mesh^10^ and IMPPAT 2.0.^9^ Comparatively, the secondary metabolite space of medicinal fungi remains much less explored. To this end, we previously created a library of secondary metabolites produced by medicinal fungi, MeFSAT^15^, via an extensive manual curation of published literature. In brief, MeFSAT compiles curated information on 149 medicinal fungi, 1830 secondary metabolites, and 184 therapeutic uses of medicinal fungi. Importantly, some medicinal fungi including mushrooms have found uses as nutraceuticals, health supplements, and superfoods for promoting healthy lifestyle.^16^ Thus, the MeFSAT library of secondary metabolites produced by medicinal fungi is an attractive chemical space which can be used in HTS for drug discovery and wellness research. Therefore, it is important to analyse the structural diversity of the MeFSAT library and compare it with other natural product libraries, approved drugs and commercial libraries.

In cheminformatics literature, several methods have been proposed for quantifying the structural diversity of chemical libraries. Previously, Medina-Franco *et al*.^17^ have proposed a systematic analysis of the scaffold diversity using cyclic system retrieval (CSR) curves and Shannon entropy. Later González-Medina *et al*.^18^ have developed the consensus diversity plot (CDP) to assess the global diversity of the chemical libraries. Subsequently, these methods have been extensively used to compare and assess the structural diversity of chemical libraries including natural products.^19–24^ Specifically, González-Medina *et al*.^21^ have done a comparative analysis of the scaffold diversity of 223 fungal secondary metabolites with approved drugs and commercial libraries. They found that the fungal secondary metabolites are structurally diverse with unique scaffolds not found in other libraries analysed by them. However, we found no previous study analysing the scaffold diversity of the secondary metabolite space specific to medicinal fungi.

In this study, we performed a systematic analysis of the scaffold diversity of the secondary metabolite space of medicinal fungi compiled in MeFSAT database. Moreover, we compared the secondary metabolite space of medicinal fungi with the scaffold diversity of 9 different chemical libraries, including natural products, approved drugs and commercial semi-synthetic libraries (Table 1). Further, we used MACCS keys structural fingerprints and molecular properties to analyse the structural diversity and diversity in terms of molecular properties, respectively, of the chemical libraries. We also used CDP to assess the global diversity of the chemical libraries. Finally, we used Generative Topographic Mapping (GTM) and Principal Component Analysis (PCA) to visualize and compare the chemical space of MeFSAT and other chemical libraries considered here.

**Table 1.** List of chemical libraries analysed in this study. For each chemical library, the table provides the number of unique chemicals and the literature reference.

| Chemical Library | Description | Number of unique chemicals | Reference |
| --- | --- | --- | --- |
| MeFSAT | Secondary metabolites of medicinal fungi | 1829 | Vivek-Ananth *et al.*, 2019<sup>15</sup> |
| Approved drugs | Approved drugs from DrugBank | 2466 | Wishart *et al.*, 2017<sup>25</sup> |
| TCM-Mesh | Phytochemicals of Chinese herbs | 10127 | Zhang *et al.*, 2017<sup>10</sup> |
| IMPPAT 2.0 | Phytochemicals of Indian medicinal plants | 17915 | Vivek-Ananth *et al.*, 2022<sup>9</sup> |
| CMAUP | Phytochemicals of medicinal and edible plants across the globe | 47187 | Zeng *et al.*, 2019<sup>12</sup> |
| NPATLAS-Bacteria | Natural products in NPATLAS of bacterial origin | 12505 | Van Santen *et al.*, 2019<sup>11</sup> |
| NPATLAS-Fungi | Natural products in NPATLAS of fungal origin | 19966 | Van Santen *et al.*, 2019<sup>11</sup> |
| MEGx | Natural product library from a commercial vendor | 6458 | https://ac-discovery.com/ |
| NATx | Semi-synthetic library from a commercial vendor | 33000 | https://ac-discovery.com/ |
| MACROx | Semi-synthetic library from a commercial vendor | 4306 | https://ac-discovery.com/ |

## Methods

### Compilation and preprocessing of chemical libraries

For this comparative analysis, the list of secondary metabolites of medicinal fungi was obtained from our previously published database, Medicinal Fungi Secondary Metabolites And Therapeutics^15^ (MeFSAT; https://cb.imsc.res.in/mefsat). The chemical diversity of the secondary metabolite space of medicinal fungi was compared with the list of approved drugs compiled in DrugBank version 5.1.9^25^, phytochemicals, microbial natural products, and commercial semi-synthetic libraries. Specifically, we considered the following phytochemical libraries namely, TCM-Mesh^10^ which compiles phytochemicals from Chinese herbs, IMPPAT 2.0^9^ which compiles phytochemicals from Indian medicinal plants, and CMAUP^12^ which compiles phytochemicals from medicinal and edible plants across the globe. Moreover, we subdivided the microbial natural product library, NPATLAS^11^, for this analysis into chemicals of fungal origin (NPATLAS-Fungi) and chemicals of bacterial origin (NPATLAS-Bacteria). Lastly, we also considered another natural product library MEGx, and two semi-synthetic libraries namely, NATx and MACROx, from a commercial vendor (https://ac-discovery.com).

Table 1 provides a summary of the different chemical libraries analysed here. Notably, the chemical libraries in SDF file format were cleaned and deduplicated to create non-redundant lists using MayaChemTools.^26^

### Computation of molecular scaffolds

The scaffolds capture the core molecular framework of a chemical, and this concept has been widely used to assess and compare the scaffold diversity of chemical libraries.^9,21,27–30^ In this study, we used the scaffold definition proposed by Bemis-Murcko^31^ to compute the molecular scaffolds of the chemicals in different libraries, wherein the scaffold is represented by all the ring systems and linkers connecting them. Based on this definition, only chemicals with cyclic systems have a scaffold. Since the analysed libraries contain both cyclic and acyclic chemicals, the acyclic chemicals have been assigned a pseudo-scaffold in this work.

Following Lipkus *et al*.^23,29^, one can compute the molecular scaffolds in different chemical libraries at three different levels namely, graph/node/bond (G/N/B) level, graph/node (G/N) level or graph level. The scaffold at G/N/B level has connectivity, element and bond information, and thus, making it more informative than G/N or graph level. Hence, we analysed the scaffold diversity of different libraries using molecular scaffolds computed at the G/N/B level for each chemical in this study. The scaffold computations were performed using custom in-house python scripts employing RDKit (https://www.rdkit.org/).

### Quantifying the scaffold diversity

Previous investigations^9,17,27,30^ have shown that cyclic system retrieval (CSR) plots help in quantifying the scaffold diversity of chemical libraries. Using the scaffold information at the G/N/B level, we plotted the CSR^28^ curves for each chemical library considered here. In a CSR curve for a chemical library, the percentage of scaffolds is plotted on the *x*-axis and the percentage of compounds containing those scaffolds is plotted on the *y*-axis. From the CSR curves, we computed two metrics namely, the area under the curve (AUC) and the percentage of scaffolds required to retrieve 50% of the chemicals (P_50_), to quantify and compare the scaffold diversity of the different chemical libraries. The scaffold diversity of a chemical library has maximum value when the corresponding CSR curve is a diagonal line, which implies that 50% of scaffolds will retrieve 50% of the compounds in the library (P_50_) and the AUC value is 0.5.

The Shannon entropy (*SF*) is employed to characterize the distribution of chemicals among the most populated scaffolds^17,32^ in a chemical library. For a selected population of *P* chemicals and top *n* scaffolds in a library, *SE* is defined as:

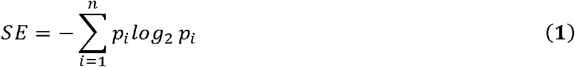

where

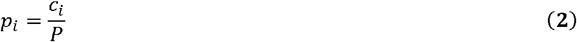

In the above equations, *c_i_* is the number of chemicals containing the scaffold *i*, and *p_i_* is the probability of the occurrence of the scaffold *i* in *P* chemicals containing a total of *n* scaffolds. The maximum possible value of *SE* is *log*_2_*n* wherein all the *P* chemicals are evenly distributed among *n* scaffolds, and this represents high scaffold diversity in the library. The minimum possible value of *SE* is 0 wherein all the *P* chemicals have the same scaffold, and this represents low scaffold diversity in the library. Since *SE* is dependent upon the number of scaffolds n, we scaled *SE* by dividing it with the maximum value of *SE*. The scaled Shannon entropy (SSE) is defined as:

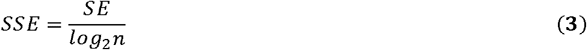

It is evident that SSE can take values from 0 to 1, where 0 corresponds to low scaffold diversity and 1 corresponds to high scaffold diversity of the chemical library.

### Inter- and Intra-library distance based on structural fingerprints and molecular properties

We quantified the inter- and intra-library distances between the different chemical libraries using structural fingerprints and molecular properties of the chemicals. We computed the Molecular ACCess System (MACCS) keys fingerprints with 166 bits for each chemical using RDKit. To compare the similarity between two libraries, we computed the Soergel distance, which is a complement of the Tanimoto coefficient, using the binary fingerprints of chemical structures.^33^ If *x* and *y* are the binary fingerprints for two chemicals then the corresponding Soergel distance can be computed as follows:

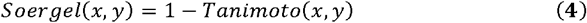

where

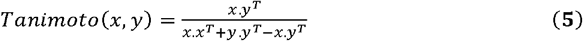

We computed the Tanimoto coefficient for a pair of chemicals using RDKit. The similarity coefficient of chemicals across two libraries, *D_u_* and *D_v_*, i.e., inter-library distance, was computed using Soergel-based inter-library distance *d_uv_* following Owen *et al*.^33^ and is given by:

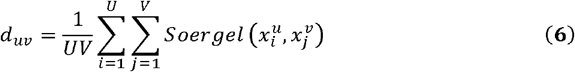

In the above equation, *U* and *V* are the number of chemicals in the two libraries *D_u_* and *D_v_*. The diversity of chemicals in a single library or intra-library distance can be computed by modifying Eq. (6) and is given by:

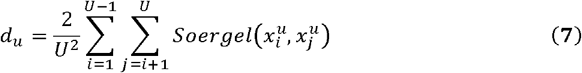

Further, we computed six molecular properties important for drug-likeness^34–36^ namely, hydrogen bond donors (HBD), hydrogen bond acceptors (HBA), octanol/water partition coefficient (LogP), molecular weight (MW), topological polar surface area (TPSA) and number of rotatable bonds (RTB), for each chemical using RDKit. Notably, these molecular properties were previously employed to compare chemical diversity across different libraries.^37^ The inter-library distance based on the six molecular properties between two libraries *D_u_* and *D_v_* containing *U* and *V* chemicals, respectively was computed by measuring the Euclidean distance function^38^ and is given by:

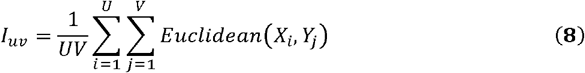

where the *Euclidean*(*X_i_, Y_i_*) is given as

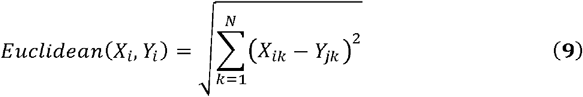

In the above equations, *X_i_* and *Y_j_* represent *N*-dimensional vectors containing molecular properties of chemicals *i* and *j* in libraries *D_u_* and *D_v_* respectively.

### Consensus diversity plots

Consensus diversity plot (CDP) is a two-dimensional visualization used to compare the diversity of chemical libraries.^18^ CDP captures four important properties to characterize the diversity of the chemical libraries. Firstly, the structural fingerprint based diversity of a library, captured by Soergel-based intra-library distance using MACCS keys fingerprints, is plotted on the x-axis of CDP. Secondly, the scaffold diversity of a library, captured by AUC from the corresponding CSR curve, is plotted on the y-axis of CDP. Thirdly, the data points in CDP are coloured using a pink to purple gradient to capture the molecular properties based intra-library distance computed using the Euclidean distance function. Fourthly, the relative size of the chemical libraries is represented by the size of the data points in CDP. Following González-Medina *et al*.^18^, we analysed the CDP by partitioning it into 4 quadrants which are differentiated by distinct colours. To define the four quadrants in CDP, we considered the median of the Soergel-based intra-library distance and AUC value of 0.75 to assign the threshold for x-axis and y-axis, respectively.

### Visualization of chemical spaces

In cheminformatics literature^33,39^, multiple methods have been proposed for dimensionality reduction and visualization of chemical spaces. Of these methods, Generative Topographic Mapping^40^ (GTM) and Principal Component Analysis^41^ (PCA) have been widely used for chemical space visualization. Using GTM and PCA, we visualized the different chemical libraries based on MACCS keys fingerprints and the six molecular properties important for drug-likeness. PCA projects the high-dimensional data to a lowdimensional space using linear mapping^41^. Although PCA is widely used for dimensionality reduction, it is unsuitable for nonlinear data^42^. In contrast, GTM is a nonlinear method that projects the highdimensional data to a two-dimensional space using Radial Basis Function^40^.

To represent any chemical space using structural fingerprints, we employed MACCS keys fingerprints with 166 binary bits that capture the presence or absence of structural features in a chemical structure. To represent any chemical space using molecular properties, we employed the six molecular properties namely, HBD, HBA, LogP, MW, TPSA and RTB, as described in a preceding section. The high-dimensional input data for a chemical library in terms of either structural fingerprints or molecular properties was then mapped to a two-dimensional space using: (a) GTM implemented using ugtm^43^ python package, and (b) PCA implemented using scikit-learn^44^ python package. Subsequently, the dataset corresponding to a chemical library after dimensionality reduction is visualized using Matplotlib^45^ python package.

## Results and discussion

### Molecular scaffolds of the secondary metabolite space of medicinal fungi

MeFSAT^15^ is a dedicated resource compiling secondary metabolites produced by medicinal fungi. After building the manually curated MeFSAT^15^ database, we had performed a detailed analysis of the chemical space captured therein. Characterization of the molecular scaffolds in a chemical library enables identification of compounds with novel scaffolds that can be considered in the drug discovery pipeline. Previously, we had not computed the molecular scaffolds for the secondary metabolites in MeFSAT^15^ database. In this study, we therefore identified the molecular scaffolds for the secondary metabolites in MeFSAT (Methods).

Next, we updated the MeFSAT database by including the valuable information on molecular scaffolds identified in each secondary metabolite at three different levels namely, G/N/B, G/N and Graph, following the definition by Lipkus *et al*.^28,29^ (Methods). In the updated MeFSAT database (https://cb.imsc.res.in/mefsat), the users can filter secondary metabolites by selecting scaffolds of interest via the ‘Scaffold filter’ tab under ‘Advanced Search’ option (SI Fig. S1). Moreover, the detailed information page for each secondary metabolite in the updated MeFSAT database now displays the identified scaffolds at the three levels (SI Fig. S1).

Overall, the secondary metabolites in MeFSAT were found to contain 618 unique scaffolds at G/N/B level including the pseudo-scaffold used to account for acyclic chemicals in the library (Table 2; Methods). Of these 618 scaffolds, 56 scaffolds occur in 5 or more secondary metabolites in MeFSAT, and Fig. 1 is a molecular cloud visualization of these frequent scaffolds after excluding the benzene ring scaffold. After computing the molecular scaffolds for the approved drug space compiled in DrugBank version 5.1.9 ^25^, we found that there is minimal overlap between scaffolds in the secondary metabolites of medicinal fungi and scaffolds in approved drugs with 94% of the scaffolds in MeFSAT not present in approved drugs (Fig. 2). This result highlights the unique scaffolds present in secondary metabolite space of medicinal fungi, and thereby, the potential of this natural product space for future drug discovery.

**Figure 1.**
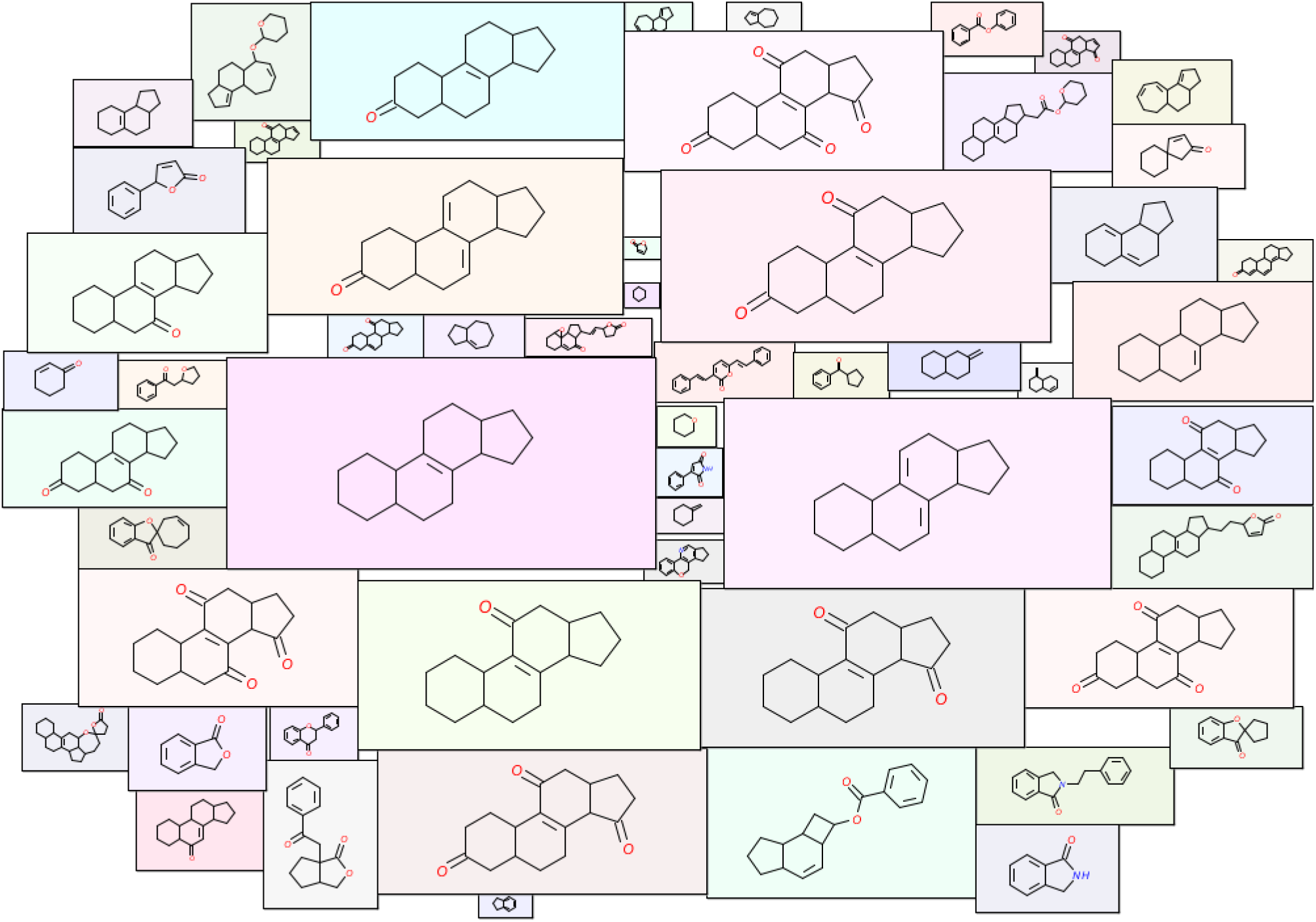
Molecular cloud visualization of the top scaffolds that occur in at least 5 secondary metabolites in MeFSAT. In this figure, the size of a scaffold image reflects its frequency of occurrence in MeFSAT. Further, we considered only the cyclic chemicals while selecting the top scaffolds. Moreover, the benzene ring scaffold is omitted from this visualization as it is the most frequent scaffold in any large chemical library.

**Figure 2.**
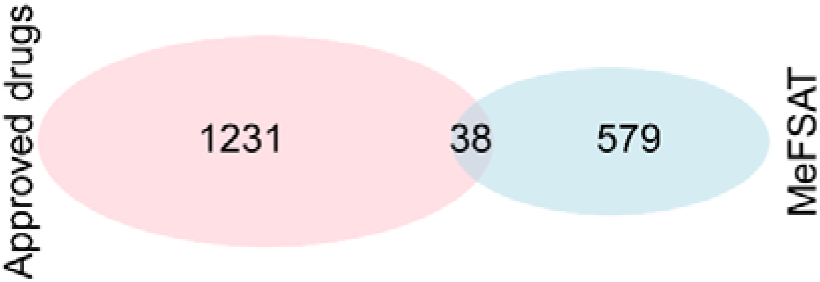
Venn diagram displays the overlap between the molecular scaffolds occurring in secondary metabolite space of MeFSAT and approved drugs in DrugBank.

**Table 2.** Comparative analysis of the scaffold diversity of the secondary metabolites in MeFSAT with other chemical libraries. Here, M is the size of the library, N is the total number of scaffolds (including the pseudoscaffold for acyclic chemicals) in the library, N_sing_ is the total number of singleton scaffolds in the library, AUC is the area under the curve for the corresponding CSR curve, and P_50_ is the percentage of scaffolds required to retrieve 50% of the chemicals in the library.

| Chemical Library | M | N | $N_{\text{sing}}$ | N/M | $N_{\text{sing}}/M$ | $N_{\text{sing}}/N$ | AUC | $P_{50}$ |
| --- | --- | --- | --- | --- | --- | --- | --- | --- |
| MeFSAT | 1829 | 618 | 370 | 0.338 | 0.202 | 0.599 | 0.786 | 7.443 |
| Approved drugs | 2466 | 1270 | 1026 | 0.515 | 0.416 | 0.808 | 0.729 | 11.102 |
| TCM-Mesh | 10127 | 3949 | 2629 | 0.39 | 0.26 | 0.666 | 0.77 | 8.787 |
| IMPPAT 2.0 | 17915 | 5184 | 3344 | 0.289 | 0.187 | 0.645 | 0.824 | 3.492 |
| CMAUP | 47187 | 11118 | 6181 | 0.236 | 0.131 | 0.556 | 0.837 | 3.913 |
| NPATLAS-Bacteria | 12505 | 4234 | 2463 | 0.339 | 0.197 | 0.582 | 0.78 | 9.258 |
| NPATLAS-Fungi | 19966 | 6414 | 3779 | 0.321 | 0.189 | 0.589 | 0.794 | 7.141 |
| MEGx | 6458 | 2566 | 1723 | 0.397 | 0.267 | 0.671 | 0.767 | 9.08 |
| NATx | 33000 | 11445 | 6370 | 0.347 | 0.193 | 0.557 | 0.764 | 11.769 |
| MACROx | 4306 | 2039 | 1329 | 0.474 | 0.309 | 0.652 | 0.719 | 16.037 |

### Comparative analysis of the scaffold diversity of secondary metabolite space of medicinal fungi with other chemical libraries

In this study, we compared the scaffold diversity of secondary metabolites of medicinal fungi compiled in MeFSAT with 9 other chemical libraries (Table 1; Methods). Table 2 provides the statistics on the number of scaffolds (N), the fraction of scaffolds per molecule (N/M), and the number of singleton scaffolds (N_sing_) for the 10 chemical libraries analysed here.

In terms of the fraction of scaffolds per molecule, the secondary metabolite space of MeFSAT (N/M = 0.338) is similar to the libraries of natural products from fungi (NPATLAS-Fungi; N/M = 0.321) and natural products from bacteria (NPATLAS-Bacteria; N/M = 0.339). Although the library of approved drugs from DrugBank and the semi-synthetic library MACROx are among the smallest in terms of library size, the two chemical libraries were found to have higher N/M ratio of 0.515 and 0.474, respectively. In terms of the fraction of singleton scaffolds per molecule, the secondary metabolite space of MeFSAT (N_sing_/M = 0.202) was found to have a higher value in comparison to relatively larger natural product libraries namely, IMPPAT 2.0, CMAUP and NPATLAS-Fungi, analysed here (Table 2). Overall, in terms of the fraction of scaffolds per molecule and the fraction of singleton scaffolds per molecule, the secondary metabolite space of MeFSAT has scaffold diversity similar or higher in comparison to other natural product libraries analysed here (Table 2).

### Analysis of scaffold diversity via cyclic system retrieval curves

Inspired by previous investigations^9,17,27,30^, we computed cyclic system retrieval (CSR) curves to quantify and compare the scaffold diversity of chemical libraries (Fig. 3; Methods). From the CSR curves shown in Fig. 3, it can be seen that the secondary metabolite space of MeFSAT has higher scaffold diversity in comparison to larger natural product libraries IMPPAT 2.0 and CMAUP. Further, from the CSR curves shown in Fig. 3, we find that the scaffold diversity of the secondary metabolite space of MeFSAT is similar to natural product libraries NPATLAS-Fungi, NPATLAS-Bacteria, TCM-Mesh and MEGx. On the other hand, we find that the approved drugs from DrugBank and the semi-synthetic library MACROx have the highest scaffold diversity among the chemical libraries analysed here.

**Figure 3.**
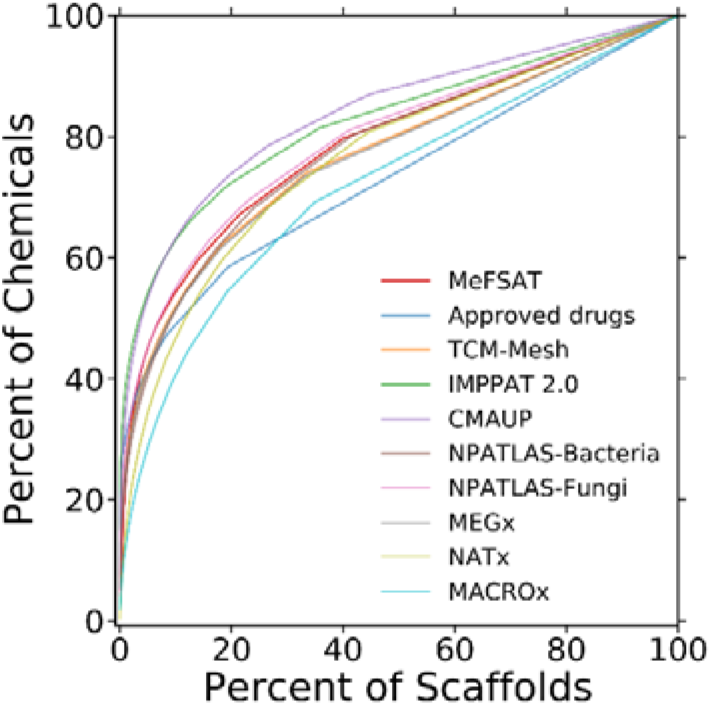
Cyclic system retrieval (CSR) curves for the 10 different chemical libraries considered in this study. Note that a CSR curve close to the diagonal line indicates high scaffold diversity. The two metrics namely, the area under the curve (AUC) and percentage of scaffolds required to retrieve 50% of the chemicals (P_50_), derived from the CSR curves also enable quantitative comparison of the scaffold diversity between chemical libraries.

Moreover, we performed a quantitative comparison of the different chemical libraries using two metrics derived from the CSR plot namely, area under the curve (AUC) and percentage of scaffolds required to retrieve 50% of the chemicals (P_50_) (Methods). As mentioned in the Methods section, a lower AUC value and a higher P_50_ value is an indicator of higher scaffold diversity. Table 2 lists the two metrics computed from CSR curves shown in Fig. 3 for the different chemical libraries analysed here. We find that the secondary metabolite space of MeFSAT has an AUC value similar to other natural product libraries. Interestingly, we also find that the P_50_ values distinguish on the one hand the semi-synthetic libraries, NATx and MACROx, and on the other hand the approved drugs from the natural product libraries analysed here (Table 2).

### Distribution of chemicals across the most populated scaffolds in different libraries

*We* computed the scaled Shannon entropy (SSE) for each chemical library to quantify the nature of distribution of chemicals across the top most populated scaffolds (Methods). The maximum value (1) of SSE indicates an even distribution of the chemicals across the top most populated scaffolds whereas the minimum value (0) of SSE indicates that all chemicals have the same scaffold. In Table 3, *we* present the computed SSE values by considering the top 5 (SSE5) to top 70 (SSE70) most populated scaffolds for each chemical library analysed here. The secondary metabolite space of MeFSAT (SSE values: 0.979 to 0.876) has the highest diversity among all the natural product libraries analysed here. The semi-synthetic libraries NATx (SSE values: 0.994 to 0.984) and MACROx (SSE values: 0.940 to 0.957) have the highest SSE values among the libraries considered here, and moreover, the SSE values are closer to 1 for the two libraries indicating high scaffold diversity. Note that the scaffold diversity interpreted from SSE values is based only on the top most populated scaffolds whereas the AUC based on CSR curves is based on analysis of all the scaffolds in a chemical library, and therefore, SSE and AUC are not the same and measure different aspects of the diversity. This explains the reason behind the approved drugs having the lowest SSE values (0.675 to 0.680) in spite of having low AUC value for the CSR curve (Fig. 3).

**Table 3.** Scaled Shannon entropy (SSE) computed using the most populated scaffolds for the chemical libraries analysed in this study. The table provides the computed SSE values for the 5 most populated scaffolds (SSE5) to the computed SSE values for the 70 most populated scaffolds (SSE70) for different chemical libraries.

| Chemical Library | SSE5 | SSE10 | SSE20 | SSE30 | SSE40 | SSE50 | SSE60 | SSE70 |
| --- | --- | --- | --- | --- | --- | --- | --- | --- |
| MeFSAT | 0.979 | 0.956 | 0.929 | 0.913 | 0.899 | 0.888 | 0.882 | 0.876 |
| Approved drugs | 0.675 | 0.618 | 0.626 | 0.64 | 0.654 | 0.663 | 0.672 | 0.68 |
| TCM-Mesh | 0.812 | 0.782 | 0.787 | 0.794 | 0.799 | 0.803 | 0.805 | 0.807 |
| IMPPAT 2.0 | 0.671 | 0.649 | 0.663 | 0.669 | 0.678 | 0.685 | 0.688 | 0.691 |
| CMAUP | 0.785 | 0.766 | 0.781 | 0.781 | 0.784 | 0.788 | 0.792 | 0.796 |
| NPATLAS-Bacteria | 0.785 | 0.766 | 0.781 | 0.781 | 0.784 | 0.788 | 0.792 | 0.796 |
| NPATLAS-Fungi | 0.849 | 0.856 | 0.863 | 0.866 | 0.867 | 0.867 | 0.867 | 0.867 |
| MEGx | 0.868 | 0.857 | 0.851 | 0.855 | 0.857 | 0.859 | 0.859 | 0.859 |
| NATx | 0.994 | 0.991 | 0.986 | 0.985 | 0.985 | 0.985 | 0.984 | 0.984 |
| MACROx | 0.94 | 0.95 | 0.952 | 0.953 | 0.955 | 0.956 | 0.957 | 0.957 |

Fig. 4 displays the distribution of the number of chemicals across the top 70 most populated scaffolds in MeFSAT, semi-synthetic library NATx, phytochemical library TCM-Mesh and the approved drugs. The corresponding distributions for other libraries analysed here are shown in SI Fig. S2. Libraries with low SSE70 value have a less even distribution of chemicals as can be seen in the case of approved drugs (Fig. 4d). In contrast, the semi-synthetic library NATx which has the highest SSE70 value has a more even distribution of chemicals (Fig. 4b).

**Figure 4.**
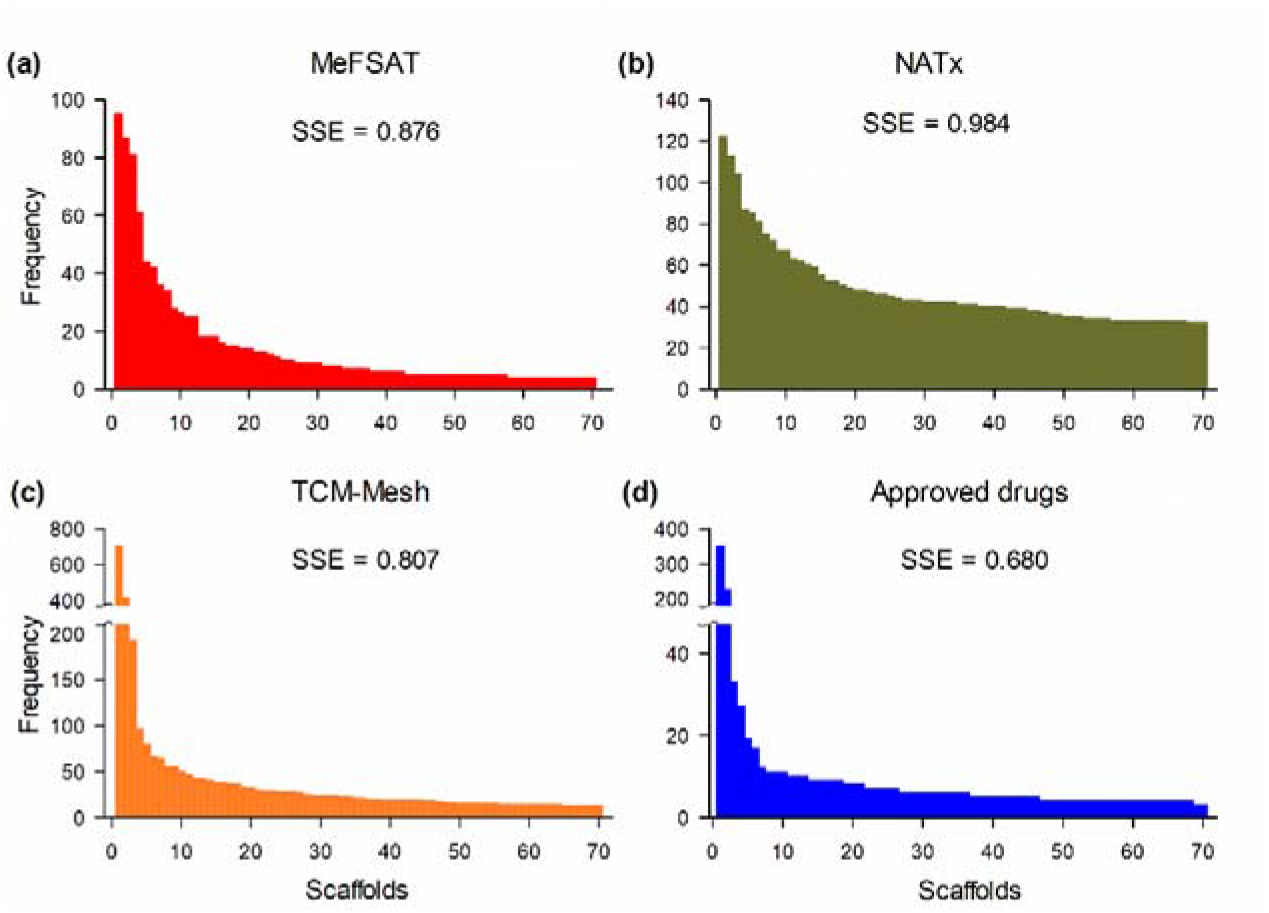
Distribution of chemicals across the top 70 most populated scaffolds in: (a) secondary metabolites in MeFSAT, (b) semi-synthetic library NATx, (c) phytochemical library TCM-Mesh, and (d) Approved drugs.

### Inter- and Intra-library distance between the secondary metabolite space of medicinal fungi and other chemical libraries

By employing the Soergel distance using MACCS keys fingerprints and the Euclidean distance using six molecular properties, we quantified the inter- and intra-library distances for the chemical libraries analysed here (Methods). Fig. 5a,b display the triangular heatmap plots (THPs) summarizing the inter- and intra-library distances for the chemical libraries based on: (a) the Soergel distance computed using MACCS keys fingerprints, and (b) Euclidean distance computed using molecular properties, respectively. In Fig. 5, the diagonal cells of THPs show the intra-library distance coloured in gradients of red, wherein darker shades of red indicate high diversity and lighter shades of red indicate low diversity. Moreover, the off-diagonal cells in THPs show the inter-library distances coloured in gradients of blue, wherein darker shades of blue indicate high inter-library distance (i.e., low similarity between the pair of libraries) and lighter shades of blue indicate low inter-library distance (i.e., high similarity between the pair of libra ries).

**Figure 5.**
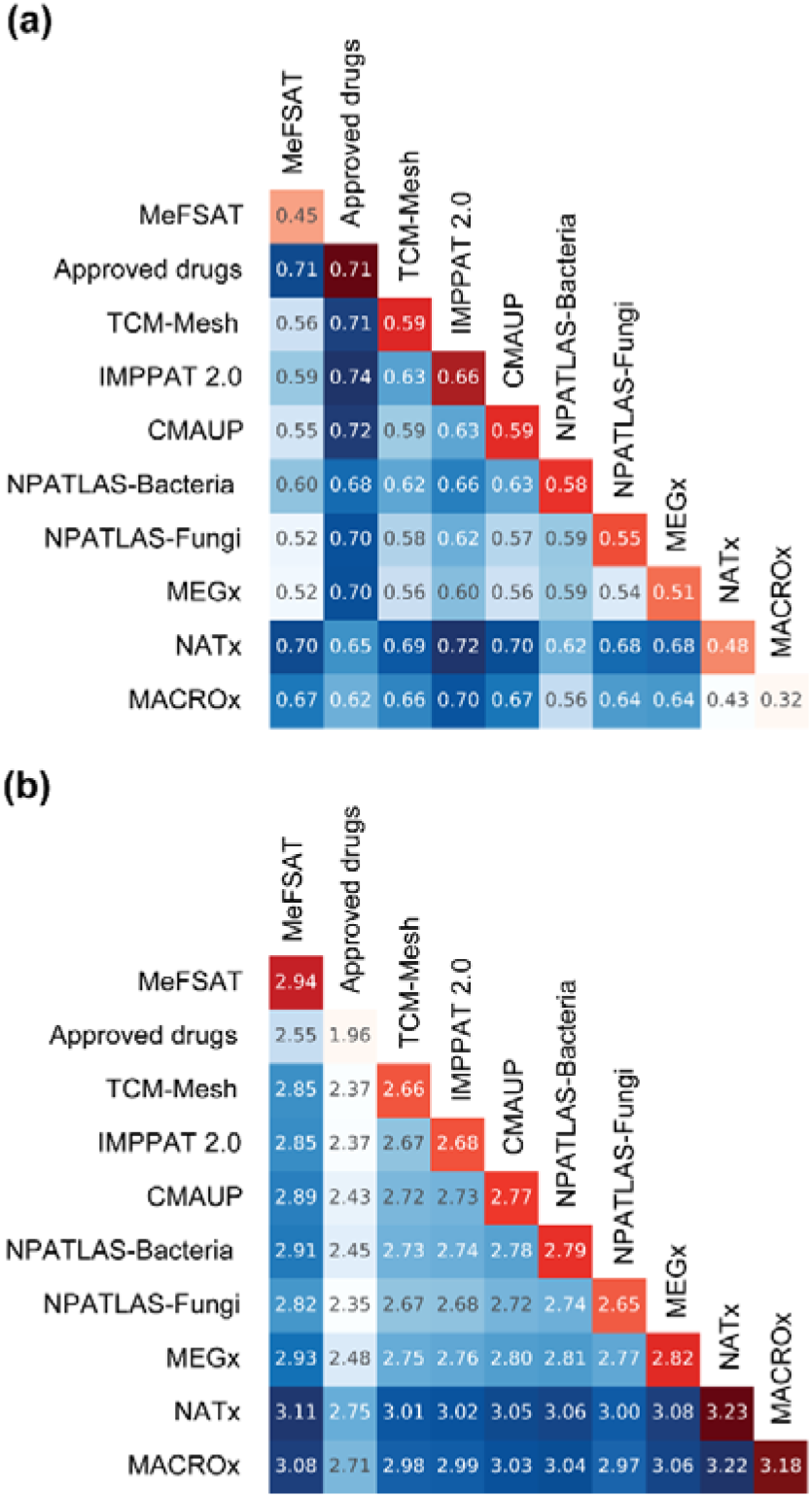
Triangular heatmap plots (THPs) for the chemical libraries analysed here, (a) THP based on Soergel distance using MACCS keys fingerprints and (b) THP based on Euclidean distance of molecular properties. The off-diagonal cells show the inter-library distance and are coloured in gradients of blue. Dark blue indicates low similarity and light blue indicates high similarity between libraries. The diagonal cells show the intra-library diversity and are coloured in gradients of red. Dark red indicates high diversity and light red indicates low diversity within the library.

#### Structural diversity based on Soergel distance using MACCS keys fingerprints

From the off-diagonal cells in THP based on structural fingerprints shown in Fig. 5a, it is evident that secondary metabolites in MeFSAT are similar to the other natural product libraries analysed here. In particular, the secondary metabolite space of MeFSAT is closest to NPATLAS-Fungi (0.52) and MEGx (0.52). In contrast, the secondary metabolite space of MeFSAT is farthest from the approved drugs (0.71) followed by semi-synthetic libraries, NATx (0.70) and MACROx (0.67). Notably, the high inter-library distance between MeFSAT and approved drugs highlights that the MeFSAT library is more suitable for HTS to identify new chemical entities. From the diagonal cells in THP based on structural fingerprints shown in Fig. 5a, it is clear that MeFSAT has an intermediate intra-library distance (0.45) whereas the approved drug space has the highest intra-library distance (0.71) followed by the IMPPAT 2.0 phytochemical space (0.66).

#### Chemical diversity based on Euclidean distance using Molecular properties

From the off-diagonal cells in THP based on molecular properties shown in Fig. 5b, it is evident that the secondary metabolites in MeFSAT are more similar to natural product libraries and approved drugs, while the secondary metabolites in MeFSAT are less similar to the semi-synthetic libraries, NATx and MACROx. In particular, the secondary metabolites in MeFSAT are closest to the NPATLAS-Fungi (2.82) based on molecular properties. Interestingly, the secondary metabolite space of MeFSAT is found to be similar to the space of approved drugs based on the molecular properties, in spite of the high inter-library distance based on structural fingerprints and minimal scaffold overlap between the two libraries. This observation highlights that the MeFSAT library is enriched with secondary metabolites with favourable molecular properties similar to approved drugs though being structurally diverse from the approved drugs, and this makes them more suitable for HTS to identify new chemical entities. From the diagonal cells in THP based on molecular properties shown in Fig. 5b, it is seen that MeFSAT has the highest intra-library distance (2.94) among the natural product libraries considered here, while the semi-synthetic libraries, NATx (3.23) and MACROx (3.18), have the highest intra-library distance across all the libraries analysed here. When comparing the structural fingerprint based intra-library distance (Fig. 5a) and the molecular properties based intra-library distance (Fig. 5b), the approved drugs were found to have the highest diversity based on structural fingerprints but low diversity based on molecular properties. This contrasting observation can be understood by the fact that the drug development pipeline is often constrained by the physicochemical properties which limit the diversity of the molecular properties of the approved drugs.^46,47^

#### Global diversity analysis with Consensus diversity plot

Fig. 6 shows the consensus diversity plot (CDP) which captures the global diversity of the chemical libraries analysed here (Methods). Briefly, in the CDP, the x-axis gives the Soergel-based intra-library distance computed using MACCS keys fingerprints, the y-axis gives the AUC from the CSR curves, the colour of the data points capture the molecular properties based intra-library distance computed using Euclidean distance function, and the relative size of the chemical libraries is reflected in the size of the data points (Methods). Furthermore, the data points (corresponding to different chemical libraries) fall in either one of the four quadrants of the CDP. In a CDP plot, the chemical libraries in quadrant IV (salmon-red) are more diverse based on both scaffold and structural fingerprints, the libraries in quadrant III (yellow) have high scaffold diversity, the libraries in quadrant I (cyan) have high structural diversity, and libraries in quadrant II (white) have relatively lower diversity (Fig. 6).

**Figure 6.**
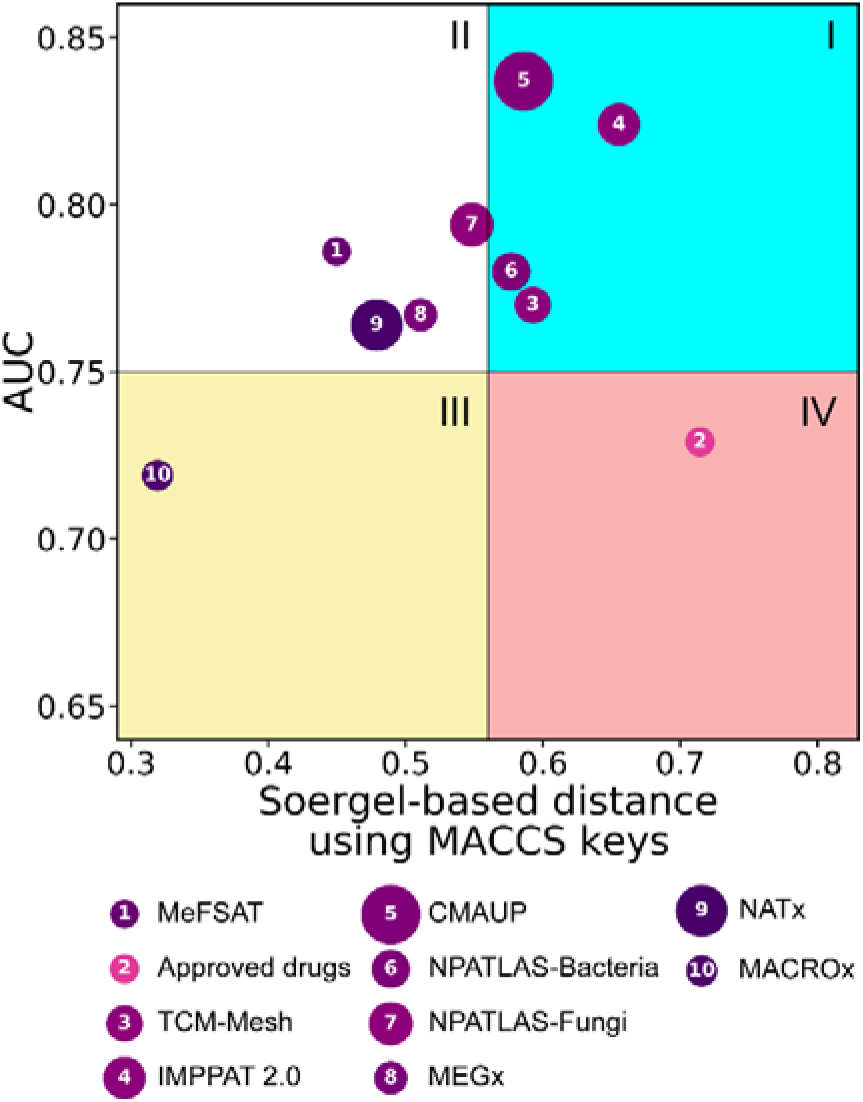
Consensus diversity plot (CDP) visualizing the global diversity of the chemical libraries. The x-axis represents the Soergel-based distance using MACCS keys and the y-axis represents the AUC from CSR curve. The CDP is divided into four quadrants: I in cyan, II in white, III in yellow and IV in salmon-red. The data points are coloured in pink to purple gradient with light pink indicating low diversity and dark purple indicating high diversity based on molecular properties. The relative size of the chemical libraries is reflected in the size of the data points.

From Fig. 6, we find that secondary metabolites in MeFSAT have higher scaffold diversity compared to larger natural product libraries such as CMAUP, IMPPAT 2.0 and NPATLAS-Fungi analysed here. Further, the secondary metabolites in MeFSAT have intermediate structural diversity similar to NPATLAS-Fungi, MEGx and semi-synthetic library NATx. Based on the colour of the data points, we find that the secondary metabolite space of MeFSAT has a similar diversity in terms of molecular properties to other natural product libraries analysed here. Moreover, we find that MeFSAT and NPATLAS-Fungi libraries are in the same quadrant of the CDP, and thus, the two libraries have similar global diversity even though the library size of NPATLAS-Fungi is ~10-fold larger than MeFSAT (Table 1). As expected, the library of approved drugs falls in the quadrant IV, underscoring the high diversity of the approved drug space. We also find that majority of the natural product libraries are in quadrant I, and thus, have high structural diversity.

By comparing the colours of the data points in Fig. 6, we find that all natural product libraries analysed here have an intermediate diversity in terms of molecular properties, whereas the semi-synthetic libraries, NATx and MACROx, have high diversity in terms of molecular properties. As can be seen in Fig. 5b, we also find that the library of approved drugs has a lower diversity in terms of molecular properties. In sum, the CDP captures the global diversity of the chemical libraries, enabling combined visual interpretations of the several metrics computed in this investigation.

### Visualization of chemical spaces

Fig. 7 is a visualization of the chemical spaces corresponding to the different libraries analysed here and the visualization was generated via GTM using MACCS keys structural fingerprints (Methods). The chemical space of the secondary metabolites in MeFSAT overlaps with the chemical space of other natural product libraries (Fig. 7), and in particular, it is found to be similar to the chemical space of NPATLAS-Fungi as per expectation (Fig. 7b). This finding also corroborates our similar observation from Fig. 5a. The chemical space of the approved drugs was found to be more spread out with minimal overlap with MeFSAT (Fig. 7b). This is in alignment with our previous findings that the secondary metabolite space of MeFSAT is structurally diverse from the space of approved drugs (Fig. 2, Fig. 5a). The chemical space of the semi-synthetic libraries, NATx and MACROx, were found to occupy a different region in the GTM based visualization which is underrepresented by the natural product libraries including MeFSAT (Fig. 7b).

**Figure 7.**
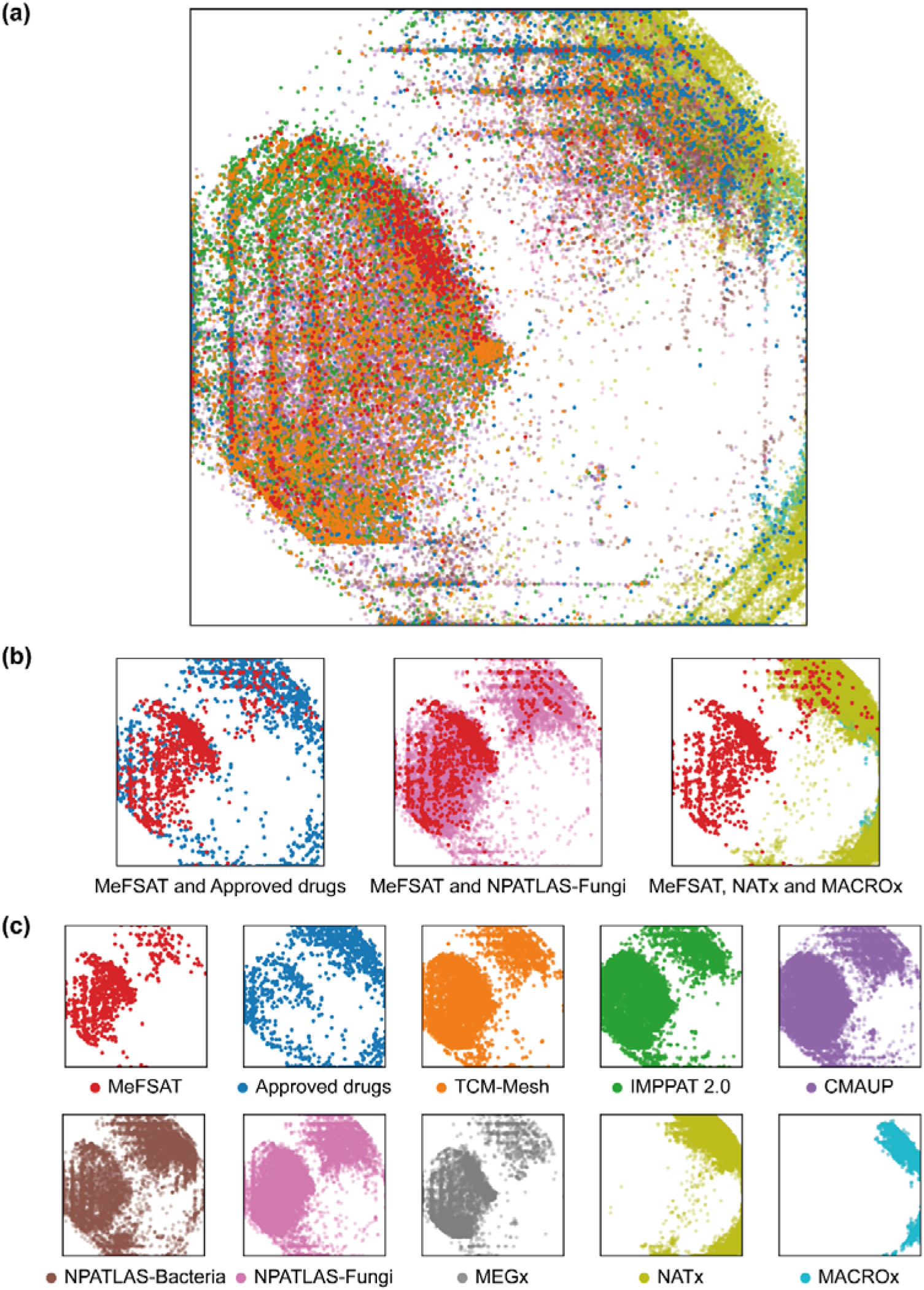
Visualization of the chemical spaces generated via GTM using MACCS keys structural fingerprints for the libraries analysed here, (a) Visualization of all chemical libraries analysed here, (b) Visualization of MeFSAT and Approved drugs, MeFSAT and NPATLAS-Fungi, and NPATLAS, NATx and MACROx. (c) Visualization of each individual chemical library. The colour used to represent each chemical library in the visualization is provided in part (c) along with the corresponding library name.

SI Fig. S3 displays visualization of the chemical spaces generated via GTM using the six molecular properties for the different libraries analysed here (Methods). The secondary metabolite space of MeFSAT is more spread out in comparison to the space of approved drugs (SI Fig. S3b). Also, the natural product libraries analysed here occupy similar regions of the chemical space wherein they occupy most regions in the two-dimensional visualization except for the regions closer to the left and bottom boundaries (SI Fig. S3c). The semi-synthetic library NATx was found to be more spread out covering regions not occupied by the natural product libraries (SI Fig. S3).

SI Fig. S4 displays visualization of the chemical spaces generated via PCA using MACCS keys structural fingerprints for the different libraries analysed here (Methods). The observations on different chemical spaces analysed here from SI Fig. S4 generated via PCA closely follow those obtained from visualization generated via GTM using MACCS keys fingerprints. The visualization of the chemical spaces generated via PCA using six molecular properties (SI Fig. S5) for different libraries analysed here was found to be less discriminative with the different libraries occupying a similar region in the lower-dimensional space.

## Conclusions

The present investigation aims to analyse and compare the scaffold and structural diversity of the secondary metabolite space of medicinal fungi (as compiled in the MeFSAT database) with nine different chemical libraries including natural products, approved drugs and semi-synthetic libraries. Firstly, we updated the MeFSAT database (https://cb.imsc.res.in/mefsat/) with the information on identified scaffolds in the secondary metabolites of medicinal fungi (SI Fig. S1).

Secondly, we compared the scaffold diversity of the secondary metabolites in MeFSAT with other chemical libraries using CSR curves. We find that the secondary metabolite space of MeFSAT has equal or higher scaffold diversity in comparison to other natural product libraries based on the fraction of scaffolds per molecule and diversity analysis using the two metrics, AUC and P_50_, computed from CSR curves (Tables 1 and 2; Fig. 3). Thirdly, based on the scaled Shannon entropy (SSE), we find that the chemicals containing the top most populated scaffolds in the MeFSAT library are more evenly distributed in comparison to other natural product libraries, and hence, the MeFSAT library is more diverse based on SSE analysis (Fig. 4; SI Fig. S2).

Fourthly, apart from analysing the scaffold diversity of the chemical libraries, we also analysed the structural diversity and diversity in terms of molecular properties with Soergel distance using MACCS keys fingerprints and Euclidean distance using six molecular properties, respectively (Fig. 5). Based on the Soergel-based inter-library distances, MeFSAT is found to be closer to other natural product libraries, and is farthest from the approved drugs and semi-synthetic libraries. In terms of molecular properties based inter-library distances, MeFSAT is closer to the natural product libraries and approved drugs, whereas it is farther from the semi-synthetic libraries. Specifically, we find that the MeFSAT library has minimal scaffold overlap with the approved drugs, and is structurally also farthest from the approved drugs, yet the secondary metabolites in MeFSAT library are closer to the approved drugs in terms of the molecular properties. This highlights the suitability of the MeFSAT library for HTS to identify new chemical entities.

Fifthly, we assessed the global diversity of the chemical libraries using CDP (Fig. 6), and found that MeFSAT has intermediate structural diversity similar to natural product libraries such as NPATLAS-Fungi and MEGx, and the semi-synthetic library NATx, and has more scaffold diversity in comparison to large sized natural product libraries such as CMAUP and IMPPAT 2.0. Further, we find that the MeFSAT and NPATLAS-Fungi fall in the same quadrant of the CDP, and thus, they have similar global diversity (Fig. 6). Sixthly, by visualizing the chemical spaces corresponding to the different chemical libraries using GTM, we find that the secondary metabolite space of MeFSAT is similar to other natural product libraries, and moreover, the secondary metabolite space of MeFSAT has minimal overlap with the approved drug space (Fig. 7).

In sum, we systematically analysed here the diversity of the secondary metabolite space of medicinal fungi as compiled in MeFSAT using molecular scaffolds, structural fingerprints and molecular properties. Overall, the results from our extensive analysis attest the diversity of the secondary metabolite space of medicinal fungi, and this renders MeFSAT as an attractive chemical space for HTS in natural product based drug discovery.

## Author Contributions

R.P.V., A.K.S., S.P.B. and A.S. designed research. R.P.V. performed the computations. R.P.V., A.K.S., S.P.B. and A.S. analysed results. R.P.V., A.K.S., S.P.B. and A.S. wrote the manuscript. A.S. conceived and supervised the project. All the authors have read and approved the manuscript.

## Acknowledgements

We thank Gokul Balaji Dhanakoti for computational support and Nagamani Sukumar for discussions. Areejit Samal acknowledges funding from the Department of Atomic Energy (DAE), Government of India, and the Max Planck Society, Germany through the award of a Max Planck Partner Group in Mathematical Biology. The funders have no role in study design, data collection, data analysis, manuscript preparation or decision to publish.

## Supplementary Information

**Figure S1.** Screenshots of the (a) Scaffold filter tab under the Advanced Search option in the updated MeFSAT database to filter secondary metabolites by selecting scaffolds of interest, and (b) the detailed information page for a secondary metabolite in the updated MeFSAT database displaying the identified scaffolds for the secondary metabolite. MeFSAT is accessible at: https://cb.imsc.res.in/mefsat.

**Figure S2.** Distribution of chemicals across the top 70 most populated scaffold in libraries: (a) MACROx, (b) NPATLAS-Fungi, (c) MEGx, (d) NPATLAS-Bacteria, (e) CMAUP, and (f) IMPPAT 2.0.

**Figure S3.** Visualization of the chemical spaces generated via GTM using molecular properties for the libraries analysed here. (a) Visualization of all chemical libraries analysed here. (b) Visualization of MeFSAT and Approved drugs, MeFSAT and NPATLAS-Fungi, and NPATLAS, NATx and MACROx. (c) Visualization of each individual chemical library. The colour used to represent each chemical library in the visualization is provided in part (c) along with the corresponding library name.

**Figure S4.** Visualization of the chemical spaces generated via PCA using MACCS keys structural fingerprints for the libraries analysed here. (a) Visualization of all chemical libraries analysed here. (b) Visualization of MeFSAT and Approved drugs, MeFSAT and NPATLAS-Fungi, and NPATLAS, NATx and MACROx. (c) Visualization of each individual chemical library. The colour used to represent each chemical library in the visualization is provided in part (c) along with the corresponding library name.

**Figure S5.** Visualization of the chemical spaces generated via PCA using molecular properties for the libraries analysed here. (a) Visualization of all chemical libraries analysed here. (b) Visualization of MeFSAT and Approved drugs, MeFSAT and NPATLAS-Fungi, and NPATLAS, NATx and MACROx. (c) Visualization of each individual chemical library. The colour used to represent each chemical library in the visualization is provided in part (c) along with the corresponding library name.

