## Supplementary material for "Scaffold and structural diversity of the secondary metabolite space of medicinal fungi": SI

### Supplementary Information (SI)

### Figures S1-S5

### for

### Scaffold and structural diversity of the secondary metabolite space of medicinal fungi

R. P. Vivek-Ananth[a,b], Ajaya Kumar Sahoo[a,b], Shanmuga Priya Baskaran[a,b], Areejit Samal[a,b,*]

[a] *The Institute of Mathematical Sciences (IMSc), Chennai 600113, India*

[b] *Homi Bhabha National Institute (HBNI), Mumbai 400094, India*

(a)

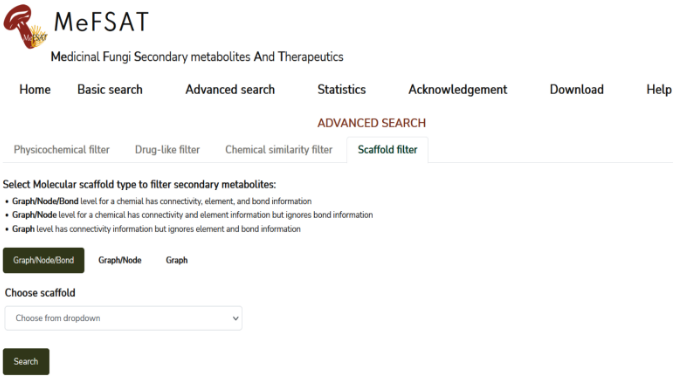


(b)

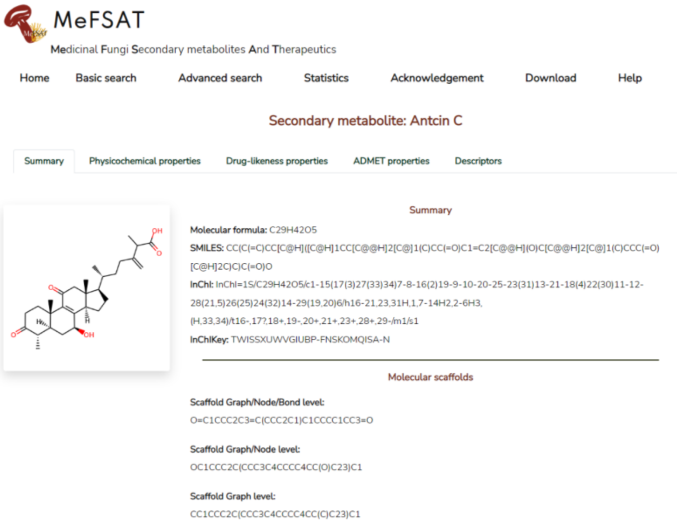


**Figure S1**: Screenshots of the **(a)** Scaffold filter tab under the Advanced Search option in the updated MeFSAT database to filter secondary metabolites by selecting scaffolds of interest, and **(b)** the detailed information page for a secondary metabolite in the updated MeFSAT database displaying the identified scaffolds for the secondary metabolite. MeFSAT is accessible at: [https://cb.imsc.res.in/mefsat](https://cb.imsc.res.in/mefsat).

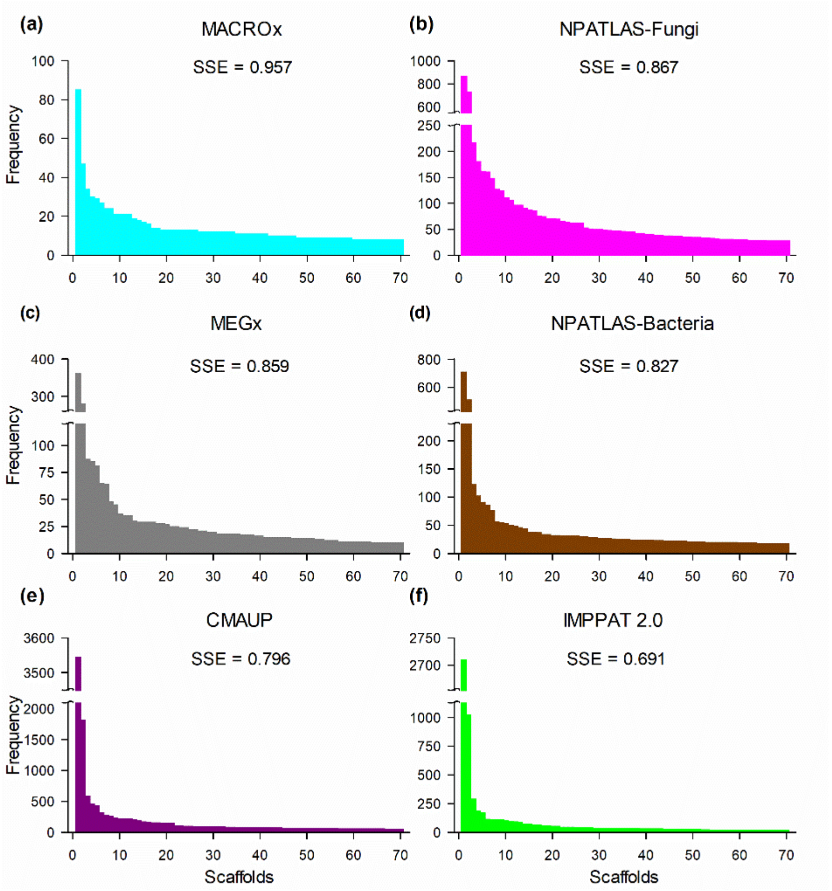


**Figure S2**: Distribution of chemicals across the top 70 most populated scaffold in libraries: (a) MACROx, (b) NPATLAS-Fungi, (c) MEGx, (d) NPATLAS-Bacteria, (e) CMAUP, and (f) IMPPAT 2.0.

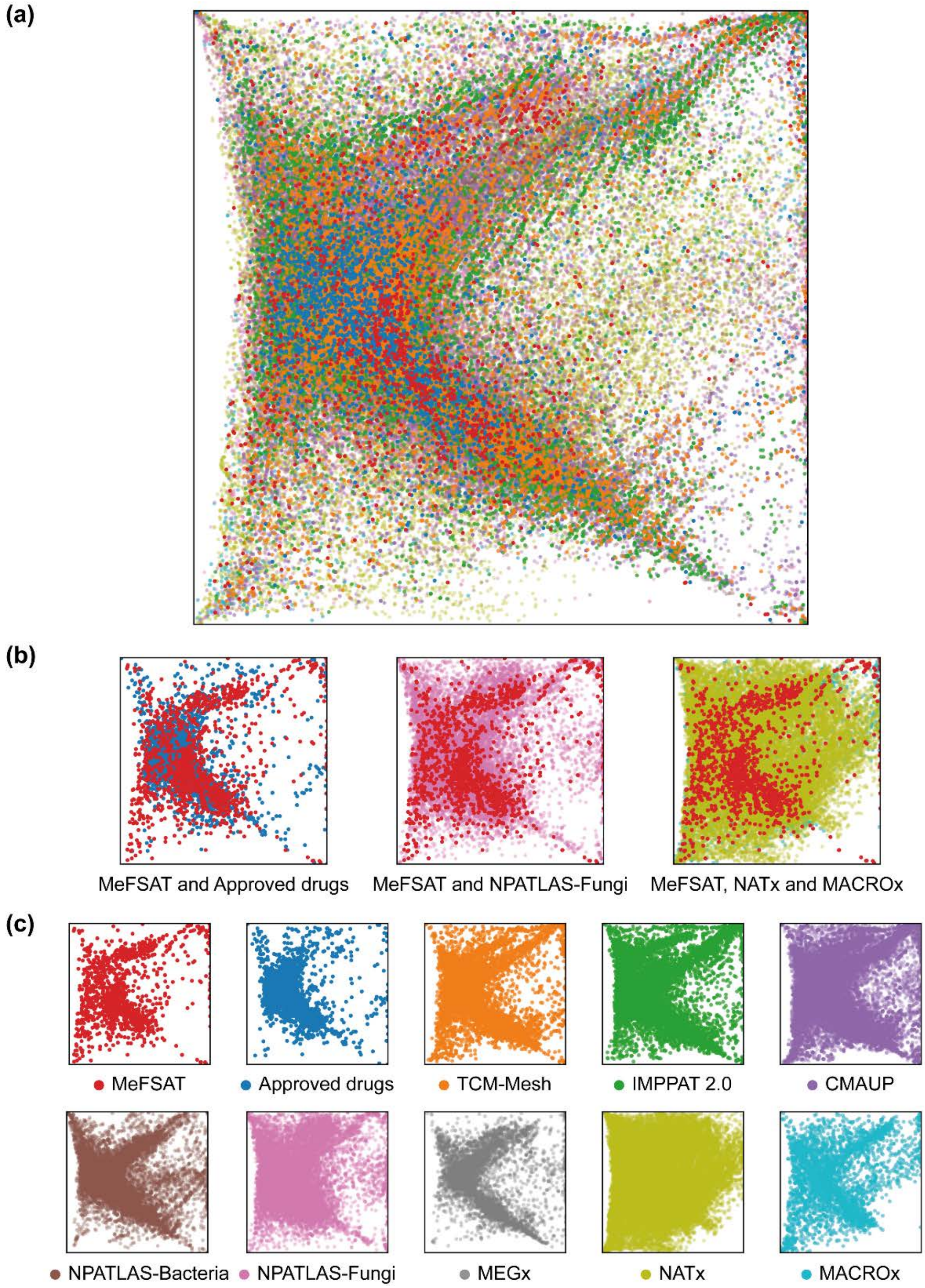


**Figure S3**: Visualization of the chemical spaces generated via GTM using molecular properties for the libraries analysed here. (a) Visualization of all chemical libraries analysed here. (b) Visualization of MeFSAT and Approved drugs, MeFSAT and NPATLAS-Fungi, and NPATLAS, NATx and MACROx. (c) Visualization of each individual chemical library. The colour used to represent each chemical library in the visualization is provided in part (c) along with the corresponding library name.

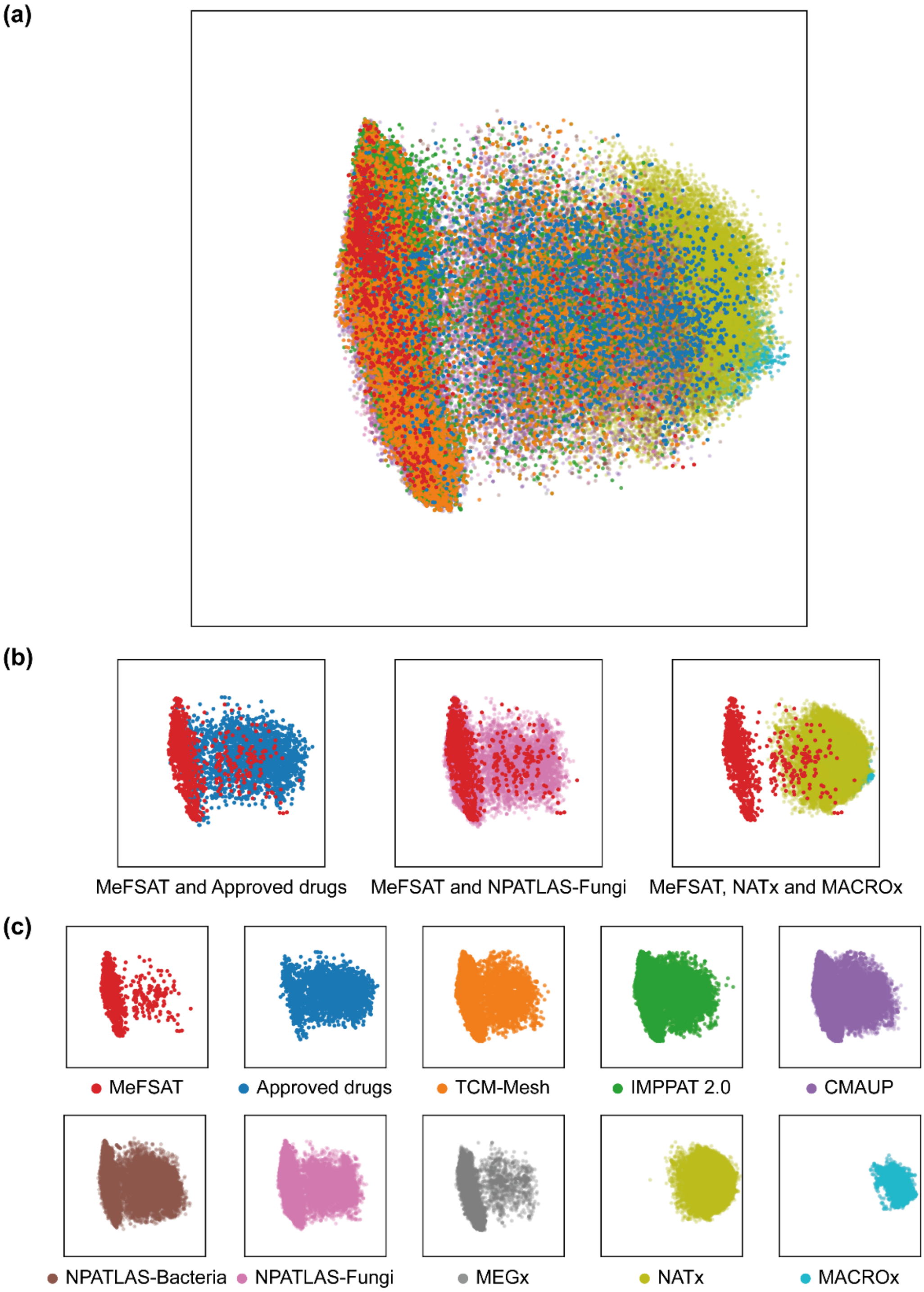


**Figure S4**: Visualization of the chemical spaces generated via PCA using MACCS keys structural fingerprints for the libraries analysed here. (a) Visualization of all chemical libraries analysed here. (b) Visualization of MeFSAT and Approved drugs, MeFSAT and NPATLAS-Fungi, and NPATLAS, NATx and MACROx. (c) Visualization of each individual chemical library. The colour used to represent each chemical library in the visualization is provided in part (c) along with the corresponding library name.

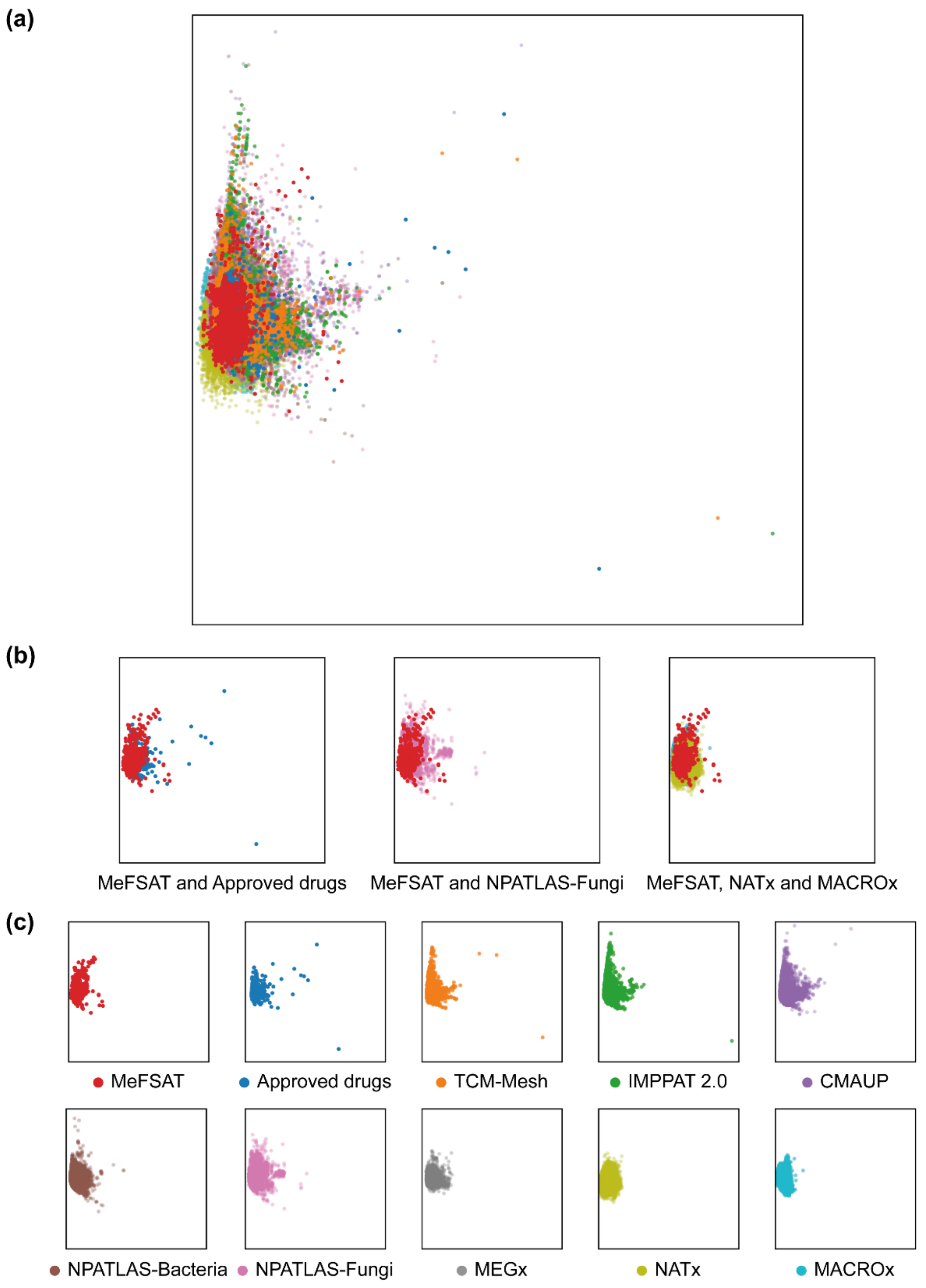


**Figure S5**: Visualization of the chemical spaces generated via PCA using molecular properties for the libraries analysed here. (a) Visualization of all chemical libraries analysed here. (b) Visualization of MeFSAT and Approved drugs, MeFSAT and NPATLAS-Fungi, and NPATLAS, NATx and MACROx. (c) Visualization of each individual chemical library. The colour used to represent each chemical library in the visualization is provided in part (c) along with the corresponding library name.
